# Microbiome composition comparison in oral and atherosclerotic plaque from patients with and without periodontitis

**DOI:** 10.1101/505982

**Authors:** Daichi Isoshima, Keisuke Yamashiro, Kazuyuki Matsunaga, Makoto Taniguchi, Takehiro Matsubara, Shuta Tomida, Kazuhiro Omori, Tadashi Yamamoto, Shogo Takashiba

## Abstract

There is no conclusive evidence regarding a causal relationship between periodontitis and atherosclerosis. In this study, we examined the microbiome in the oral cavity and atheromatous plaques from atherosclerosis patients with or without periodontitis to investigate the role of oral bacteria in the formation of atheromatous plaques. We chose four patients with and without periodontitis, who had undergone carotid endarterectomy. Bacterial samples were extracted from saliva on the tongue surface, from periodontal pocket (during the oral examination), and from the atheromatous plaques. We investigated the general and oral conditions from each patient and performed next-generation sequencing analysis for all bacterial samples. There were no significant differences between both groups concerning general conditions. However, the microbiome patterns of the gingival pocket showed differences depending on the absence or presence of periodontitis, while those of the saliva were relatively similar. The microbiome pattern of the atheromatous plaques was entirely different from that in saliva present on the tongue surface and gingival pocket, and oral bacteria were seldom detected. However, the microbiome pattern in atheromatous plaques was different in the presence or absence of periodontitis. These results indicated that oral bacteria did not affect the formation of atheromatous plaques directly. However, the metabolic products of microbiome or the host inflammatory response might indirectly influence the composition of atheromatous plaques.

## Introduction

More than 100 trillion microbes reside in niches within the human body. Collectively, these microbes constitute the microbiome [1]. The co-existence and interactions between eukaryotic and microbial cells is vital in regulating physiological functions. The microbiome balance has been linked with obesity, cancer, intestinal disorders, and mental disorders [2, 3], and periodontitis [4].

Periodontitis is a predominant oral infectious disease in which an excessive immune response directed at the microbiome on the tooth surface destroys the periodontal tissue, forming periodontal pockets. The microbiome in this pocket includes pathogenic anaerobic bacteria that can form biofilms, which are inherently a drug-resistant and challenge host immunity [5]. The mature biofilm causes further periodontitis progression because of the prolonged inflammation associated with the protracted immune response. Therefore, periodontitis has two main features; it is an infectious disease caused by microbiome imbalance and a chronic inflammatory disease caused by a dysregulated immune response.

These two characteristic features of periodontitis are shared by various systematic diseases, including diabetes, arteriosclerosis, cardiovascular diseases, brain diseases, cancer, and non-alcoholic steatohepatitis and with preterm low birth weight [6–11]. We also reported a case where the microbiome with pathogenic periodontal bacteria was implicated as the cause of infective endocarditis [12]. Arteriosclerosis includes atherosclerosis, in which an atherosclerotic plaque is formed on the blood vessel walls through various mechanisms [13]. An association between atherosclerosis and periodontitis has been suggested by some epidemiology reports. Moreover, bacterial investigations aimed at detecting periodontal bacteria in the atherosclerotic plaque have been conducted [14–16]. Although observational data support an association between periodontitis and atherosclerotic vascular disease, the data do not yet justify a causative relationship [17]. Multiple common factors, such as diabetes, high blood pressure, dyslipidemia, and smoking, affect disease progression, and there is little data of a direct involvement of periodontal bacteria in the development of atherosclerotic vascular disease [18, 19].

An association between periodontitis and atherosclerotic vascular disease was demonstrated *in vivo* using *Porphyromonas gingivalis* [20]. A clinical analysis sought to detect the DNA of periodontal bacteria in atherosclerotic plaques [21]. The impact of the disruption of the normal microbiome on various diseases is unclear and a comprehensive analysis is necessary, since the microbiome could contribute to the formation of microbiome atheroma.

Microbiome analysis has typically involved bacterial culture [22]. However, recent comprehensive bacterial analyses using the gene for 16S ribosomal RNA (rRNA) have been successful in detecting bacteria that are difficult to cultivate [23]. Next-generation sequencing (NGS) has become a popular means of examining a large number and volume of samples [24]. In this study, the association between periodontitis and atherosclerosis in the context of the microbiome in the oral cavity and atherosclerotic plaque was investigated by NGS.

## Materials and Methods

### Ethics statement

This study was approved by the ethics committee of Okayama University Graduate School of Medicine, Dentistry, and Pharmaceutical Sciences and Okayama University Hospital (Authorization Number: 1603–059) and Brain Attack Center Ota Memorial Hospital (Authorization Number: 121). All enrolled patients provided written informed consent for the use of their resected tissue and oral samples.

### Participants

The study focused on 12 patients who visited Brain Attack Center Ota Memorial Hospital between April 2016 and March 2018, and who were diagnosed with internal carotid artery stenosis. The patients were ≥ 40 years of age, underwent carotid endarterectomy, had more than ten teeth, and consented to participate.

### Samples

Atheromatous plaques (AP) were extracted from an internal carotid artery. Bacteria in the gingival pocket (GP) were collected using absorbent paper points (United Dental Manufactures Inc., Johnson City, TN, USA). Bacteria in saliva from the tongue surface (ST) were collected using forensic swabs (Sarstedt AG & Co. Nümbrecht, Germany). Blood was collected from each patient and serum prepared as previously described [25].

### Oral examination

Periodontal examinations were performed to evaluate the average pocket probing depth (PPD) and rate of bleeding on probing (BOP) for each teeth of each patient. The patients were then divided into three groups according to the Japanese Society for Periodontology Clinical Practice Guideline for the Periodontal Treatment: periodontally healthy (control, n = 4; H1-H4), mild periodontitis (n = 4, excluded from further analysis), and severe periodontitis (periodontitis, n = 4; P1-P4).

### DNA purification

APs were extensively minced using a scalpel and suspended in phosphate buffered saline (PBS). The collected material from paper points and swabs were resuspended using PBS. One milliliter of each resuspended bacterial sample was transferred to 2 ml Lysing Matrix B tubes (MP Biomedicals, Santa Ana, CA, USA) containing 0.1 mm silica beads and 500 μl ATL buffer (Qiagen, Hilden, Germany). The contents of each tube were homogenized using FastPrep 24 (MP Biomedicals) for 45 s at 6.5 m/s. Bacterial DNA was extracted using the QIAamp DNA Microbiome Kit (Qiagen) according to the manufacturer’s instructions. The quality and quantity of the DNA were verified using the NanoDrop 2000 spectrophotometer (Thermo Fisher Scientific, Wilmington, DE, USA) and the PicoGreen dsDNA assay kit (Life Technologies, Grand Island, NY, USA).

### Polymerase chain reaction and NGS analysis

The first polymerase chain reaction (PCR) using 16S rRNA primers (forward: 5’-AGAGTTTGATCCTGGCTCAG-3’, reverse: 5’-CGGTGTGTACAAGGCCCGGGAACG-3’) and KAPA HiFi HotStart ReadyMix (Kapa Biosystems Inc., Wilmington, MA, USA). Thermal cycling conditions were as follows: heating at 98°C for 3 min; 25 cycles of 98°C for 30 s, 55°C for 30 s, and 72°C for 30 s; and a final extension at 72°C for 5 min. A second PCR was performed using the first PCR amplicons and V3-V4 primers (forward: 5’-TCGTCGGCAGCGTCAGATGTGTATAAGAGACAGCCTACGGGNGGCWGCAG-3’, reverse: 5’-GTCTCGTGGGCTCGGAGATGTGTATAAGAGACAGGACTACHVGGGTATCTAATCC-3’) using the same reaction conditions. The quality and quantity of the DNA were verified using Qubit 4 Fluorometer (Invitrogen, Life Technologies, Grand Island, NY, USA) and the D1000 ScreenTape system (Agilent Technologies, Santa Clara, CA, USA).

NGS was conducted using the MiSeq^®^ system (Illumina Inc., San Diego, CA, USA). The obtained sequence was compared to the database using the CLC Genomics Workbench (CLC bio, Aarhus, Denmark). Principal component analysis (PCA) and clustering analysis were performed using R statistical software [26]. We also performed co-occurrence analysis for the 13 highly detected operational taxonomic units from control and periodontitis samples using the Quantitative Insights Into Microbial Ecology approach [27].

### Plasma IgG antibody titer test against periodontal bacteria

Plasma IgG antibody titer against periodontal bacteria was determined as described previously [28]. The selected periodontal pathogenic bacteria were *Aggregatibacter actinomycetemcomitans* (Aa) Y4, Aa ATC29523, Aa SUNY67, *Eichenerra corrodens* (Ec) FDC1073, *Fusobacterium nucleatum* (Fn) ATCC25586, *Prevotella intermedia* (Pi) ATCC25611, Pi ATCC33563, *Capnocytophaga ochracea* (Co) S3, *Porphyromonas gingivalis* (Pg) FDC381, Pg SU63, T*reponema denticola* (Td) ATCC35405, and *Tannerella forsythia* (Tf)ATCC43037.

### General condition evaluation

General conditions of patients were evaluated based on age, disease history, body mass index, blood pressure, C-reactive protein, cholesterol, and HbA1c (Table 1).

**Table 1:**
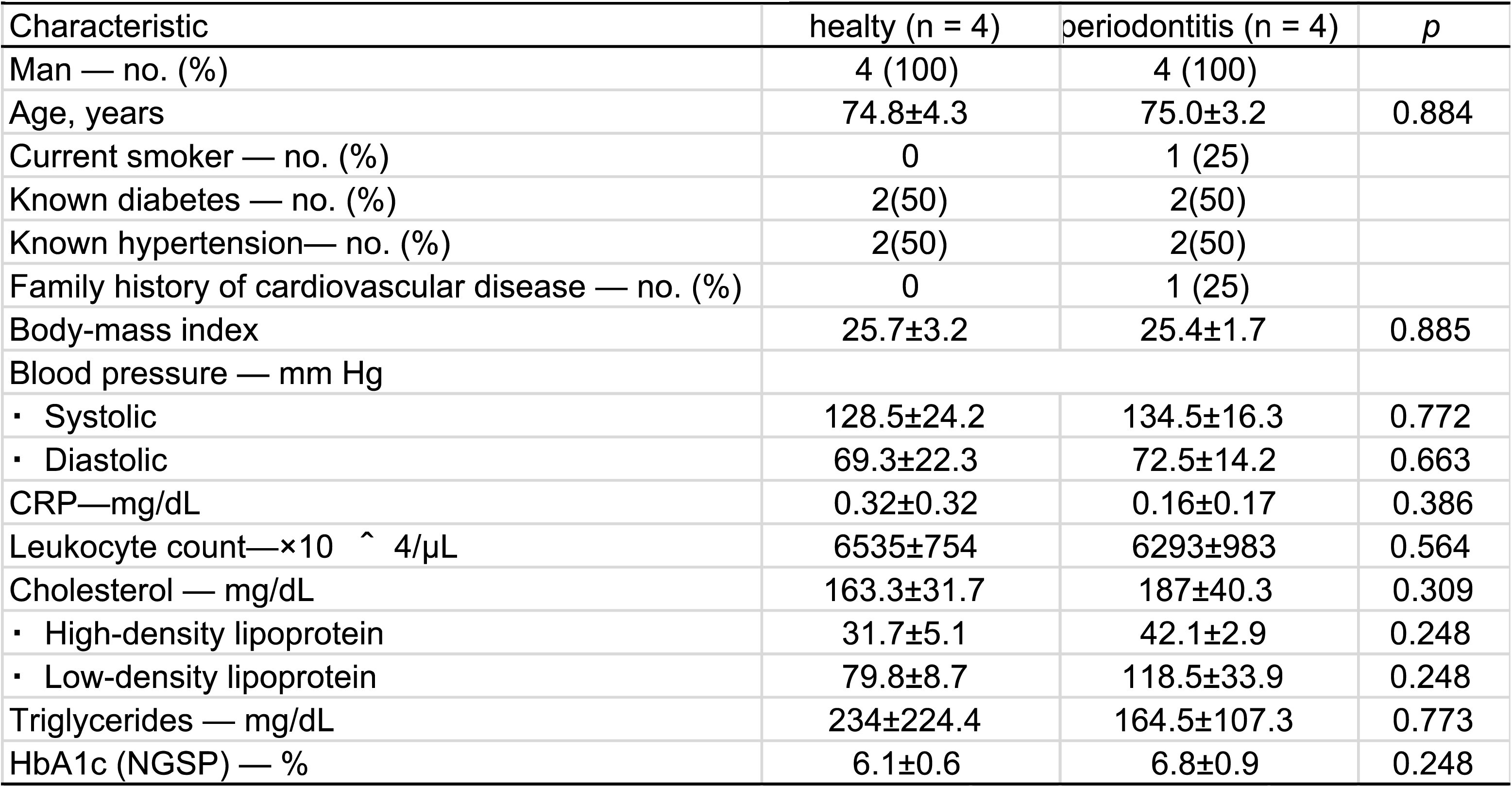
Characteristics of study participants

| Table 1 Characteristics of study participants |  |  |  |
| --- | --- | --- | --- |
| Characteristic | healty (n = 4) | periodontitis (n = 4) | p |
| Man — no. (%) | 4 (100) | 4 (100) |  |
| Age, years | 74.8±4.3 | 75.0±3.2 | 0.884 |
| Current smoker — no. (%) | 0 | 1 (25) |  |
| Known diabetes — no. (%) | 2(50) | 2(50) |  |
| Known hypertension— no. (%) | 2(50) | 2(50) |  |
| Family history of cardiovascular disease — no. (%) | 0 | 1 (25) |  |
| Body-mass index | 25.7±3.2 | 25.4±1.7 | 0.885 |
| Blood pressure — mm Hg |  |  |  |
| • Systolic | 128.5±24.2 | 134.5±16.3 | 0.772 |
| • Diastolic | 69.3±22.3 | 72.5±14.2 | 0.663 |
| CRP—mg/dL | 0.32±0.32 | 0.16±0.17 | 0.386 |
| Leukocyte count—×10<sup>4</sup>/μL | 6535±754 | 6293±983 | 0.564 |
| Cholesterol — mg/dL | 163.3±31.7 | 187±40.3 | 0.309 |
| • High-density lipoprotein | 31.7±5.1 | 42.1±2.9 | 0.248 |
| • Low-density lipoprotein | 79.8±8.7 | 118.5±33.9 | 0.248 |
| Triglycerides — mg/dL | 234±224.4 | 164.5±107.3 | 0.773 |
| HbA1c (NGSP) — % | 6.1±0.6 | 6.8±0.9 | 0.248 |

### Oral condition evaluation

We evaluated the periodontal condition for each patient group from oral examination and plasma IgG antibody titer test (Table 2).

**Table 2:** Periodontal Disease infection of study participants

| Table 2 Periodontal Disease infection of study participants |  |  |  |
| --- | --- | --- | --- |
| Variable | Controls (n = 4) | Patients (n = 4) | p |
| Total no. of teeth | 24.8 ± 2.2 | 19.8 ± 7.4 | 0.561 |
| periodontal pocket depth |  |  |  |
| • 1~3mm — ( %) | 97.9 ± 1.8 | 71.4 ± 4.5 | 0.021 |
| • 4~6mm — ( %) | 2.0 ± 1.8 | 27.2 ± 4.3 | 0.021 |
| • over 7mm — ( %) | 0.2 ± 0.3 | 2.9 ± 2.1 | 0.026 |
| Sites with gingival bleeding (%) | 7.8 ± 8.1 | 24.0 ± 12.8 | 0.081 |
| Serume IgG Antibody Titer Test against Periodontal Bacteria |  |  |  |
| • *Aggregatibacter actinomycetemcomitans* (Y4) | -0.06 ± 0.49 | 1.85 ± 1.33 | 0.021 |
| • *Aggregatibacter actinomycetemcomitans* (ATCC29523) | 0.02 ± 0.38 | 1.96 ± 0.83 | 0.021 |
| • *Aggregatibacter actinomycetemcomitans* (SUNY67) | 0.04 ± 0.71 | 2.35 ± 1.21 | 0.021 |
| • *Capnocytophaga ochracea* (S3) | -2.70 ± 2.53 | 1.94 ± 3.08 | 0.043 |
| • *Eichenerra corrodens* (FDC1073) | -0.43 ± 0.91 | 0.43 ± 1.68 | 0.564 |
| • *Fusobacterium nucleatum* (ATCC25586) | -0.11 ± 1.52 | 2.42 ± 3.82 | 0.248 |
| • *Prevotella intermedia* (ATCC33563) | -0.95 ± 1.44 | -0.63 ± 0.63 | 0.248 |
| • *Prevotella intermedia* (ATCC25611) | -0.22 ± 0.98 | 0.50 ± 1.37 | 0.773 |
| • *Porphyromonas gingivalis* (FDC 381) | -0.98 ± 0.31 | 1.83 ± 1.63 | 0.021 |
| • *Porphyromonas gingivalis* (SU 63) | -0.70 ± 0.46 | 2.58 ± 4.25 | 0.083 |
| • *Treponema denticola* (ATCC35405) | -0.03 ± 2.14 | 0.37 ± 1.94 | 0.564 |
| • *Campylobacter rectus* (ATCC33238) | 3.44± 6.40 | 17.35 ± 18.47 | 0.083 |
| • *Bacteroides forsythus* (ATCC43037) | -0.90 ± 0.25 | -0.41 ± 0.80 | 0.248 |

### Statistical analysis

The statistical analysis was performed using the Mann-Whitney *U* Test. A *P*-value of 0.05 was considered significant and was determined using SPSS Ver. 23 (SPSS Inc., Chicago, IL, USA) for all the experimental results.

## Results

The participants’ characteristics are presented in Table 1. There were no significant differences between both groups in terms of age, sex, other disease such as diabetes, and markers of inflammation and cholesterol. The periodontal disease conditions of the participants are presented in Table 2. In the control group, the ratio of PPD was < 3 mm, while the ratio of PPD in the periodontitis group was significantly higher i.e., > 4 mm. Serum IgG antibody titer was significantly higher in those with periodontitis that in control group for Aa Y4, Aa ATCC29523, Aa SUNY67, Co S3, and Pg FDC381.

### Characterization of microbiome in ST, GP, and AP

The microbiome pattern of ST was relatively similar between control samples and periodontitis samples (Fig. 1A). Among them, the ratio of *Filifactor* sp., which was reported to be virulent [29], was significantly higher in periodontitis patients than in the respective controls (Fig. 1B).

**Fig. 1.**
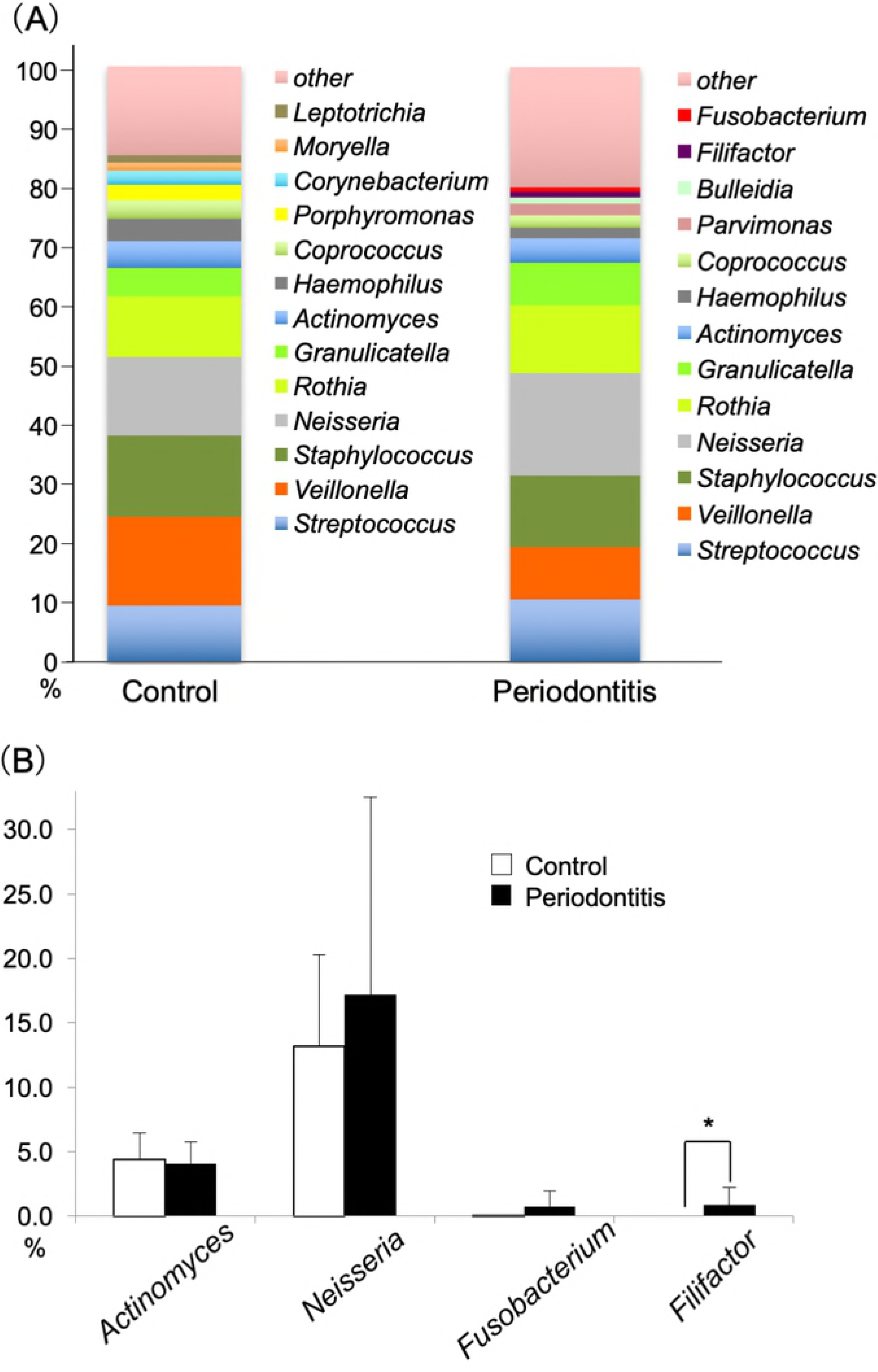
Characterization of microbiome in ST. The average ratio of the bacteria in ST from control and periodontitis patient samples is presented. (A) Bacterial genera are indicated. (B) Thirteen bacterial species were highly detected by NGS analysis. * indicates P < 0.05; Mann-Whitney *U* Test.

The microbiome pattern of GP was notably different between the control and periodontitis samples (Fig. 2A). The ratio of *Rothia* sp. and *Neisseria* sp., which exist in a healthy oral cavity, were lower in periodontitis than in control samples. Conversely, the ratios of *Fusobacterium* sp. and *Filifactor* sp., which are present in the periodontitis oral cavity, were higher in periodontitis than in control samples (Fig. 2A, B). The ratio of *Desulfobulbus* sp., which was detected in the periodontal pocket in a recent report [30], was significantly higher in periodontitis samples than in controls (Fig. 2B).

**Fig. 2.**
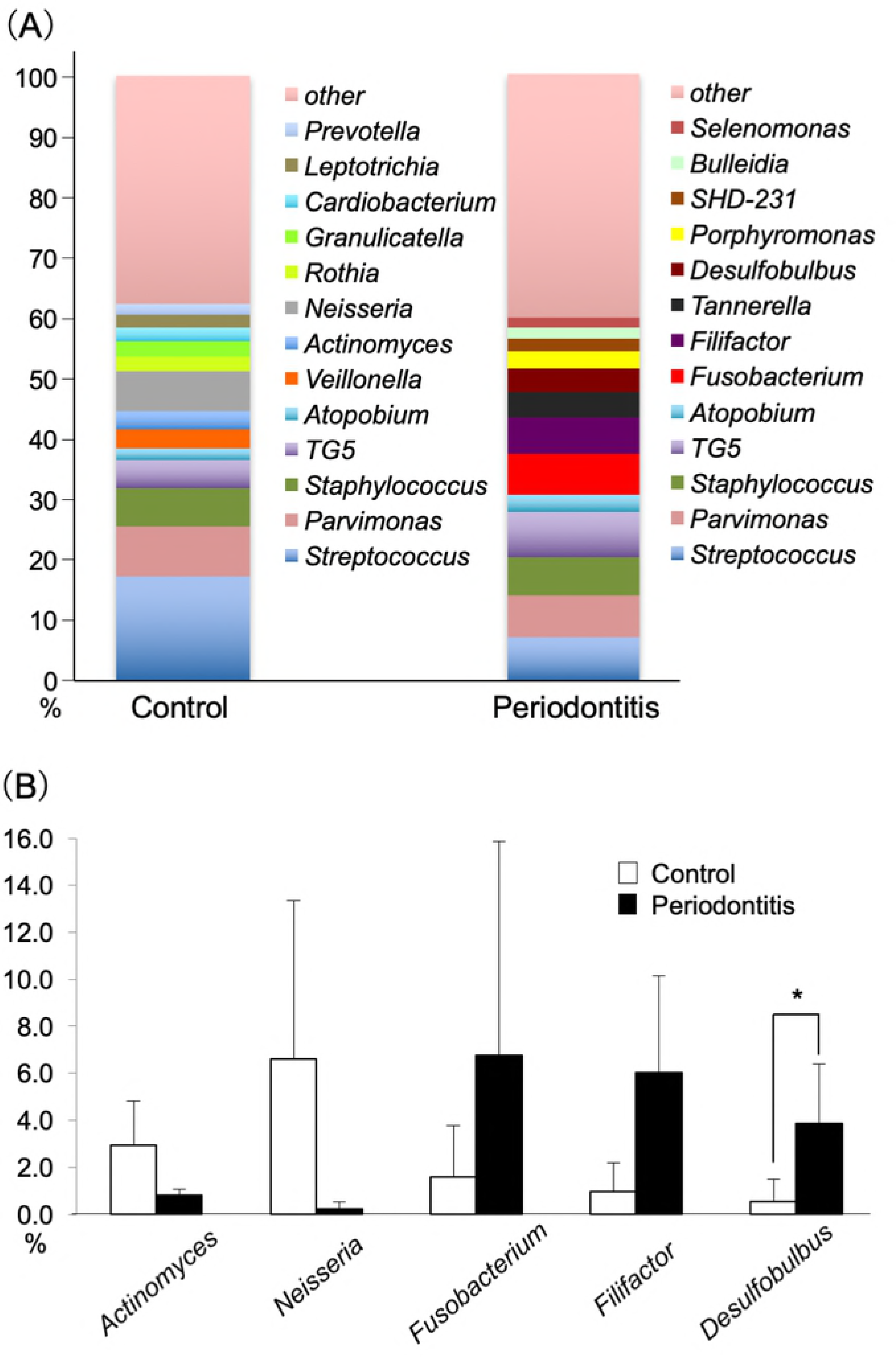
Characterization of microbiome in GP. The average ratios of the bacteria in GP from control and periodontitis samples are shown. (A) Bacterial genera are indicated. (B) Thirteen bacteria that were highly detected from the NGS analysis. * indicates P < 0.05; Mann-Whitney *U* Test.

The majority of the bacteria found in the AP microbiome belonged to the soil bacterial families *Burkholderiales, Bacillale*, and *Rhizobiales*. Their ratios were similar between periodontitis and control patients (Fig. 3). The ratio of *Sphingomonadales,* which is a normal constituent of the human bacteria flora, was higher in periodontitis samples than in control samples.

**Fig. 3.**
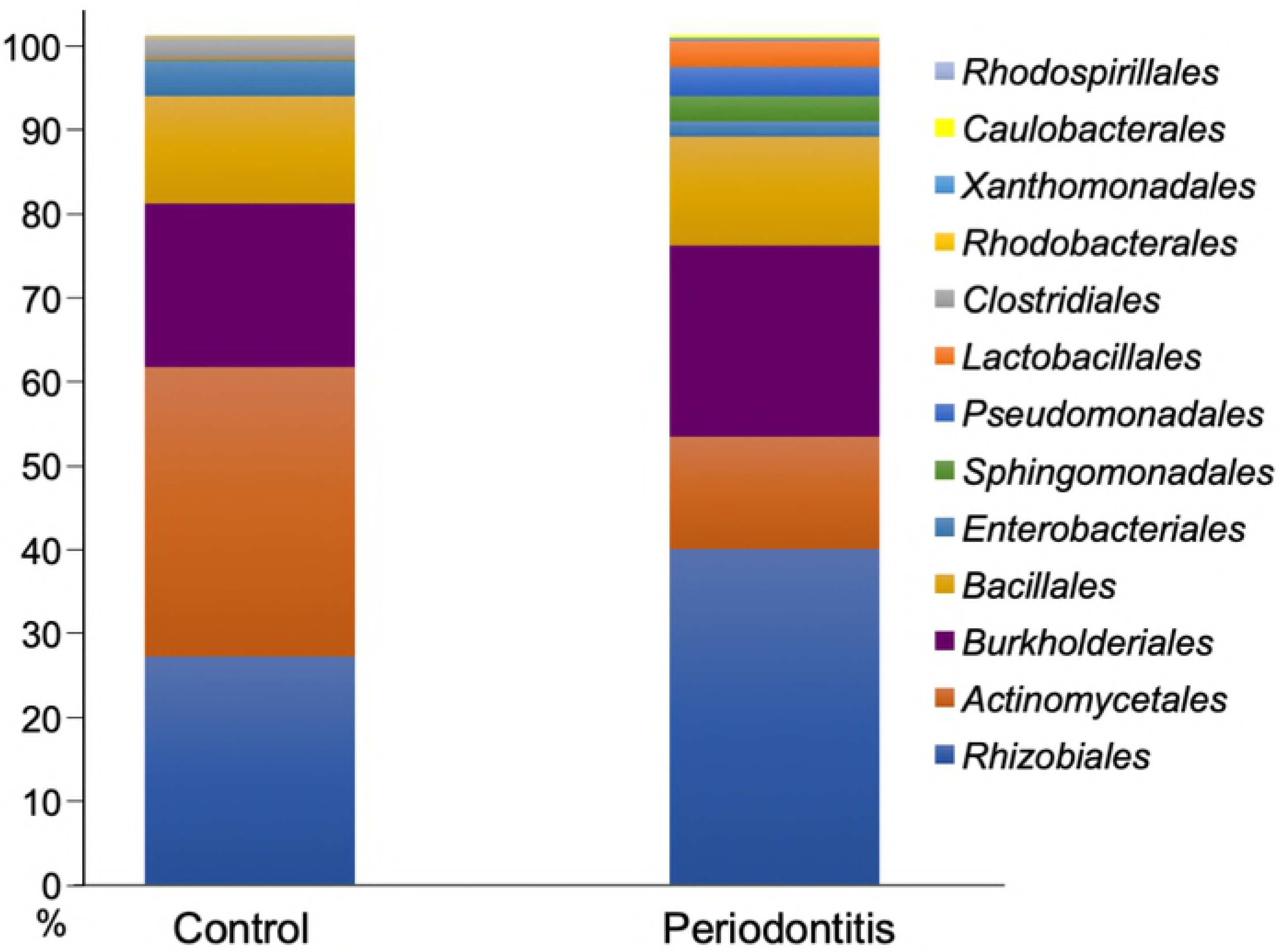
Characterization of microbiome in AP. The average ratio of the bacteria in GP from our patients (control and periodontitis samples) is depicted and the order of the bacteria is shown. Thirteen bacteria were every evident from the NGS analyses.

### Comparison of PCA results between controls and periodontitis samples

Seventy five percent of the GP bacteria from periodontitis and control samples were positioned in the center right side of the PCA graph (separated by a red solid circle) and center of the plot (separated by a blue solid circle), respectively (Fig. 4). The two circular locations were sufficiently separated. ST bacteria from periodontitis samples were located in the center left side of the panel (separated by a red dotted circle). This position was slightly more toward to the right side than that of control samples (separated by a blue dotted circle). These two circular locations were comparatively closer. The bacteria in AP were located towards the lower middle region of the plot (separated by a green solid circle), and the control and periodontitis samples could not be clearly distinguished. The AP bacteria were located far from the oral samples (ST and GP).

**Fig. 4.**
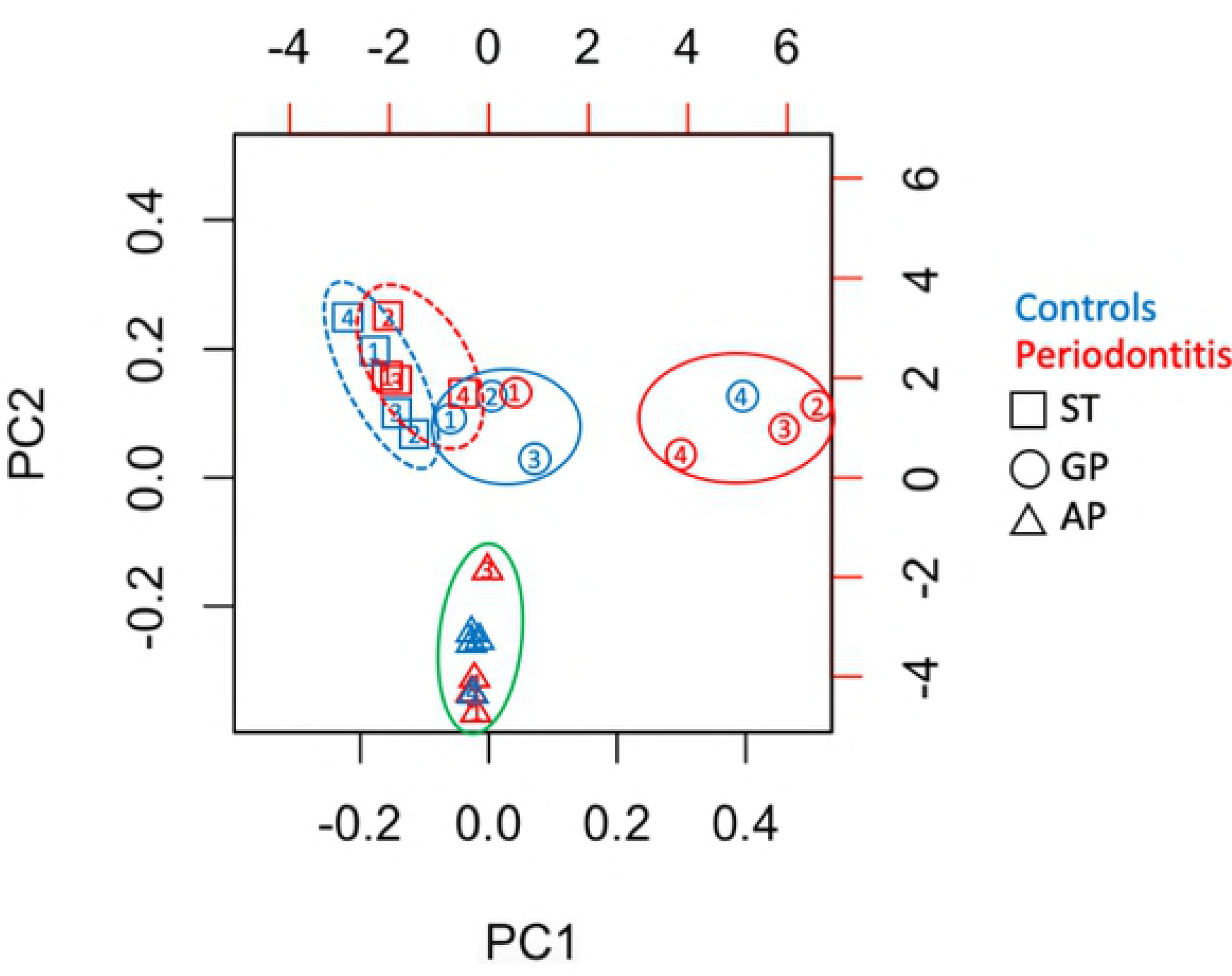
Comparison of PCA results between control and periodontitis samples. The PCA results from each sample are identified tagged as follows: □: ST; Saliva from tongue surface, ○: GP; Gingival pocket,Δ: AP; Atheromatous plaques; controls: blue color; periodontitis: red color; three GP bacteria from periodontitis and one GP from controls, blue solid circle; three GP bacteria from controls and one GP from periodontitis, red dotted circle; ST bacteria from periodontitis, blue dotted circle; ST bacteria from controls, green solid circle; AP samples from both groups.

### Comparison of clustering analysis results between control and periodontitis samples

Similar to the PCA results, bacteria from oral samples (ST and GP) and AP were completely different (Fig. 5). However, 75% of AP bacteria from periodontitis were located at the lower part in the cluster and 75% of AP bacteria from controls were located at the upper part of the periodontitis samples.

**Fig. 5.**
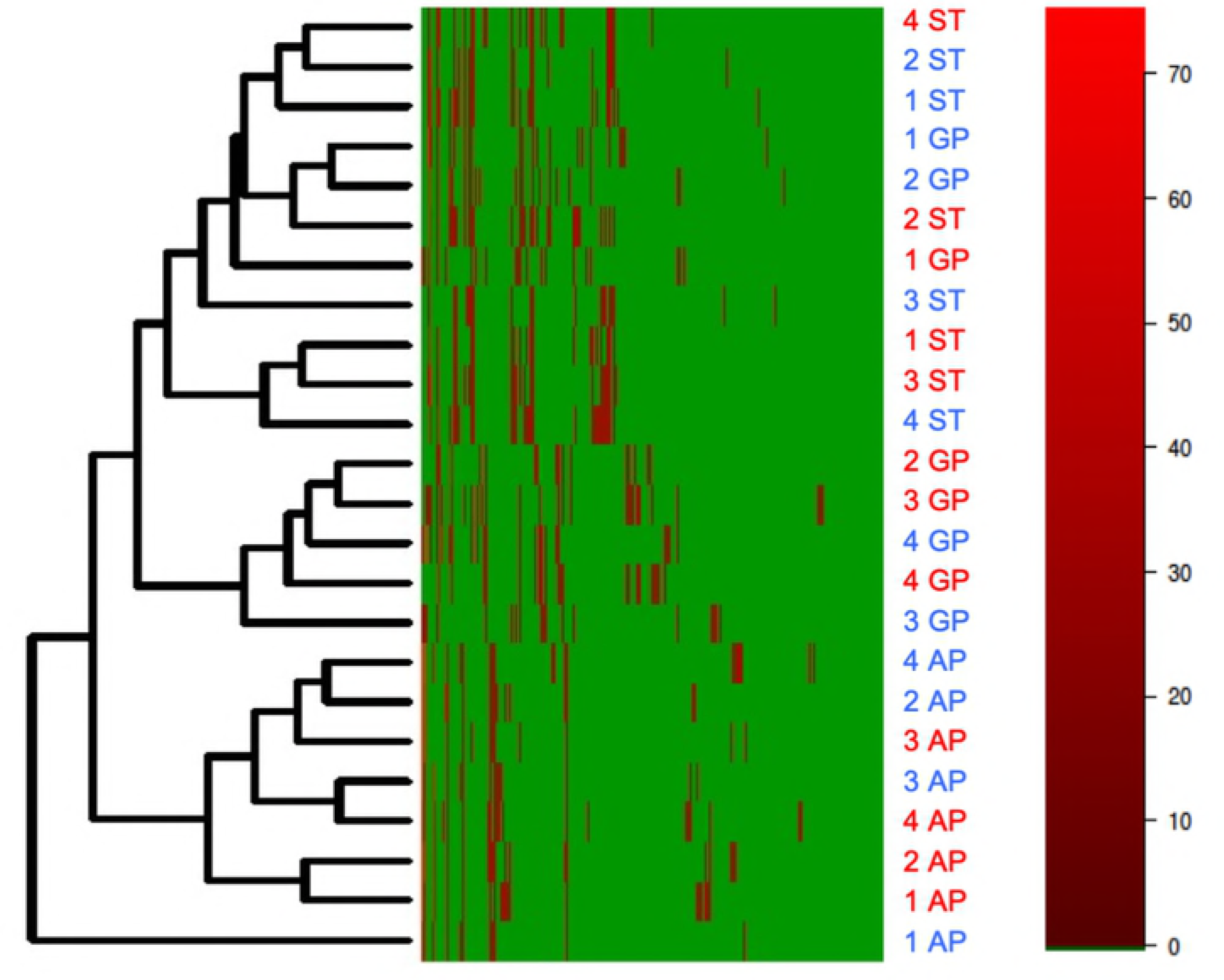
Comparison of clustering analysis results between control and periodontitis samples. The clustering analysis results from each sample were tagged as follows: ST; Saliva from tongue surface, GP; Gingival pocket, AP; Atheromatous plaques. Blue text represents control samples and red text specifies periodontitis samples.

### Co-occurrence analysis of microbiome in AP

We evaluated the correlation of the microbiome in AP between the control and periodontitis group. In both groups, the major bacteria of the network were *Agrobacterium* sp., *Delftia* sp., and *Rhizobium* sp. This echoed a previous report [26]. However, the network around *Cutibacterium acnes* was different between control and periodontitis samples (Fig. 6).

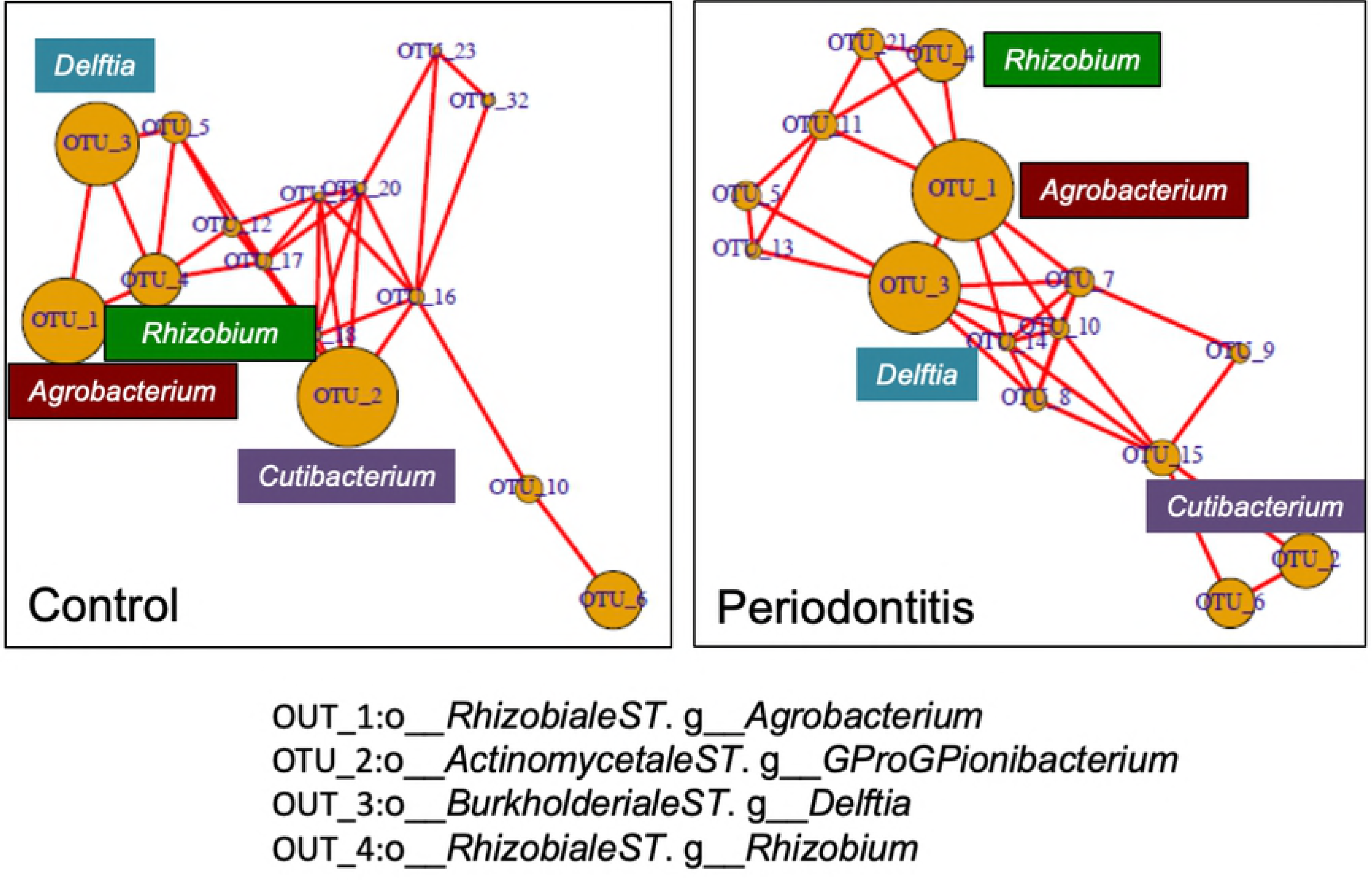

## Discussion

The human body is a complex habitat for about 1,000 species and 100-1000 trillion bacteria, wherein approximately 100 million bacteria specifically reside exclusively in the oral cavity [31]. Periodontitis is an infection caused by the members of the microbiome in the oral cavity, and is related to a number of systematic diseases. However, the detailed mechanism underlying this infection is still not completely understood [17]. As pathogenic factors for periodontitis, Red complex species (*P. gingivalis, T. denticola,* and *T. forsythia*) have been the focus of functional investigations [32]. Although there is no doubt regarding their relationship to periodontitis development, the microbiome is likely not comprised just of the pathogenic bacteria, but includes a mixture of various and diverse species of bacteria, with the total population ultimately affecting the development of this disease [33]. It has been suggested that 17 novel bacteria including *Filifactor alosis* probably induce periodontitis, even though these bacteria were not previously thought to be periodontitis-specific pathogenic bacteria [34].

The normal bacterial flora in the oral cavity, which was previously disregarded as insignificant, is actually very crucial for periodontitis development or progression. In general, pathogenic bacteria, such as *P. gingivalis*, configure the microbiome with the normal bacteria flora [29]. If the balance of pathogenic and normal bacteria in the microbiome is lost for some reason, the microbiome increases its pathogenicity and induces the disease. Therefore, a comprehensive microbiome analysis is necessary to investigate normal as well as pathogenic bacteria composition.

In this study, we performed a comprehensive microbiome analysis of the internal carotid artery stenosis in patients affected with and without periodontitis. We harvested bacteria samples from TS, GP, and AP from each patient and performed NGS analysis. This analysis showed that the microbiome in the oral cavity was more numerous for *Fusobacterium* sp. and *Filifactor* sp., which are periodontal bacterial pathogens, in the periodontitis group compared to the control group. In particular, the ratio of *Filifactor* sp. in TS was significantly higher in the periodontitis group in comparison to the control group. Conversely, the ratio of *Rothia* sp and *Neisseria* sp in GP, which are constituents of the normal bacterial flora in a healthy oral cavity, was lower in the periodontitis group than in the control group [35, 36]. Thus, remarkably, the ratio of normal bacteria in GP and TS decreased while that of pathogenic bacteria increased.

To investigate the possibility that periodontal bacteria might contribute to atheromatous plaque formation directly on the vascular wall by hematogenous spread, NGS analysis was done using the atheromatous plaque samples. Previous reports established that *P. gingivalis* induces the expression of vascular cell adhesion molecule 1 from vascular endothelial cells, and promotes thrombus formation by macrophage invasion into blood vessels, resulting in platelet aggregation [37, 38]. Another report demonstrated that *P. gingivalis* infection accelerates the progression of atherosclerosis in a heterozygous apolipoprotein E-deficient murine model [20]. Presently, oral bacteria were barely detectable in AP, regardless of the presence or absence of periodontitis. The patterns of the microbiome in AP were entirely different in TS and GP. A prior study reported detection of some oral bacteria in the atheromatous plaque [39]. However, other authors reported that *P. gingivalis* infection in an animal model induced atheromatous plaque formation, although it was actually not detected in the atheromatous plaque [40]. Our data enables us to conclude that it is unlikely that the oral bacteria spread hematogenously and directly induce the formation of atheromatous plaque on the aortic wall.

The co-occurrence analysis of the microbiome in AP revealed the most significant bacteria were *Agrobacterium* sp., *Delftia* sp., and *Rhizobium* sp., which constituted the network in both groups. Although these are soil bacteria, they were also previously detected in AP [26]. Another significant bacterium, *C. acnes,* configured the network in the control group. The relationship with *C. acnes* was different among the control and periodontitis groups. This bacterium is categorized as a normal bacterium present on the skin and in the gut, although it was also detected in AP [41]. *C. acnes* reportedly can cause sarcoidosis, sepsis, and infective endocarditis, and heat-killed *C. acnes* render mice very susceptible to lipopolysaccharide (LPS) toxicity. *C. acnes* also promote the production of cytokines, such as interleukin-12, interferon-gamma, and Toll-like receptor 4 [42]. Presently, it is conceivable that LPS produced by periodontal bacteria activated *C. acnes* in the blood vessels, which then formed the atheromatous plaque. In this scenario, the difference of the network in AP between the control and periodontitis samples might be caused by LPS that is spread hematogenously, as well as by the chronic inflammatory effect. Recently, it was reported that the production of trimethylamine-N-oxide, which promotes atherosclerosis, depends upon the metabolism of the intestinal microbiome [43]. The prior and present data indicate that the loss of microbiome balance in the human body affects the development of atherosclerosis. Periodontitis has a great effect on the microbiome configuration in the oral cavity and promotes the formation of various metabolic products. However, this detailed mechanism of atherosclerosis development remains largely unknown. In a further study, we intend to investigate the relationship between periodontitis and atherosclerosis.

## Conclusion

The ratio of oral bacteria in AP was remarkably low, and the microbiome pattern was entirely different from that found in the oral microbiome. In other words, oral bacteria did not directly induce the atheromatous plaque configuration. However, the microbiome pattern and the correlation of the microbiome in AP were different between the controls and periodontitis samples. Thus, metabolic products of the microbiome, or the host’s inflammatory response, might indirectly affect the atheromatous plaque configuration.

## Acknowledgments

We thank all the staff in the Department of Biobank, Okayama University Hospital for helping our research. We wish to acknowledge Dr. Masaru Kuriyama, Dr. Yutaka Shimoe, Dr. Sinzo Ota, Dr. Sinichi Takeshima from the Brain Attack Center Ota Memorial Hospital for collecting clinical samples, and Dr. Zulema Arias from Okayama University Graduate School of Medicine, Dentistry and Pharmaceutical Sciences for supporting this study.

